# *KNOX HOMOLOGS SHOOT MERISTEMLESS* (*STM*) and *KNAT6* are epistatic to *CLAVATA3* (*CLV3*) during embryonic meristem development in *Arabidopsis thaliana*

**DOI:** 10.1101/2020.11.11.378539

**Authors:** Sharma Nidhi, Liu Tie

## Abstract

In *Arabidopsis*, the genes *SHOOT MERISTEMLESS* (*STM*) and *CLAVATA3* (*CLV3*) antagonistically regulate shoot meristem development. *STM* is essential for both development and maintenance of the meristem, as *stm* mutants fail to develop a shoot meristem during embryogenesis. *CLV3*, on the other hand, negatively regulates meristem proliferation, and *clv3* mutants possess an enlarged shoot meristem. Genetic interaction studies revealed that *stm* and *clv3* dominantly suppress each other’s phenotypes. *STM* works in conjunction with its closely related homologue *KNOTTED1-LIKE HOMEOBOX GENE 6 (KNAT6)* to promote meristem development and organ separation, as *stm knat6* double mutants fail to form a meristem and produce a fused cotyledon. In this study, we show that *clv3* fails to promote post-embryonic meristem formation in *stm-1* background if we also remove *KNAT6. stm-1 knat6 clv3* triple mutants result in early meristem termination and produce fused cotyledons similar to *stm knat6* double mutant. Notably, the *stm-1 knat6* and *stm-1 knat6 clv3* alleles lack tissue in the presumed region of SAM. *stm knat6 clv3* also showed reduced inflorescence size and shoot apex size as compared to *clv3* single or *stm clv3* double mutants. In contrast to previously published data, these data suggest that *stm* is epistatic to *clv3* in postembryonic meristem development.

**Highlight:** *STM* and *KNAT6* genes determine post-embryonic meristem formation and activity in Arabidopsis. *clv3* mutation is unable to rescue the *stm knat6* meristemless phenotype.

## Introduction

Plants are sessile organisms and therefore have developed internal mechanisms to survive various unfavorable conditions such as drought, cold, and pathogen attack. Under mild drought conditions, plants slow their growth and produce fewer leaves to conserve resources. Unlike animals, plants have the capacity to form organs throughout their life cycle. Thus, plants maintain a reservoir of undifferentiated cells, termed meristem cells, to continuously produce organs such as leaves, stems, and flowers. This stem cell population is maintained via coordination of meristem cell proliferation with differentiation and subsequent incorporation into the organ primordium at the shoot apex. The meristem at the shoot apex, the Shoot Apical Meristem (SAM), and the meristem at the root tip, the Root Apical Meristem (RAM), are both formed during embryogenesis and are responsible for generating the aboveground and belowground organs, respectively. In *Arabidopsis*, the SAM is a roughly triangular-shaped dome, about 60 μm across at its widest point, and consists of a few hundred cells (Barton, 2010). Based on differences in morphology and cell division rates, the SAM broadly consists of two zones, the peripheral and central zones (Steeves and Sussex, 1989). The undifferentiated stem cells reside in the central zone and divide to form daughter cells, which are destined to take on organ identity and are pushed to the peripheral zone. These primordial cells will then differentiate into leaves or flowers.

Considering the importance of continuous organ production, plants have developed strategies to balance the number of self-renewing stem cells with that of differentiated cells. The meristem size and activity are tightly regulated via a delicate network of genes (Yadav et al., 2014). In *Arabidopsis, SHOOT MERISTEMLESS* (*STM*) and the *CLAVATAs* (*CLAVATA1/CLV1, CLAVATA2/CLV2*, and *CLAVATA3/CLV3*) appear to play critical roles in the regulation of shoot meristem development. We will largely focus on these two sets of antagonistic genes for the purpose of this paper.

*STM*, a *KNOTTED-Like* or *KNOX* transcription factor, is required for initiating embryonic meristem. *stm1* mutants fail to develop a meristem at the embryonic stage (Barton and Poethig, 1993). *STM* is also required to maintain the stem cell population in the SAM (Clark et al 1996). STM first appears in the globular-heart stage embryo and continues to be expressed in the SAM throughout shoot development, but is downregulated in primordial cells on the SAM periphery that differentiate into cells of the lateral vegetative organs such as leaves and bracts (Jackson et al., 1994; Long et al., 1996; Long and Barton, 2000; Hay and Tsiantis, 2010). It is this downregulation of *STM* in addition to other cues, such as auxin levels in the peripheral cells of the SAM, that triggers the initiation of organ primordia (Long and Barton, 2000; Reinhardt et al., 2000; Benková et al., 2003; and Heisler et al., 2005).

In addition to regulating meristem identity, *STM* also functions in boundary specification. *CUP-SHAPED COTYLEDON 1 (CUC1), CUC2* and *CUC3, NAC* (*NAM/ATAF/CUC*) transcription factors, control shoot organ boundary formation throughout development (Aida et al., 1997, 1999; Vroemen, 2003). At the globular stage, *CUC1* and *CUC2* are expressed in a narrow stripe separating the two developing cotyledon primordia (Aida et al, 1999; Takada et al, 2001), where they trigger *STM* expression (Long et al., 1996; Aida et al., 1999). On the other hand, during post-embryonic development, *STM* promotes expression of *CUC1/2/3*, but also represses *CUC1/2* (Kwon et al., 2006; Bosca et al., 2011; Spinelli et al., 2011). These decades of research confirm that *STM* and *CUC* regulate the SAM and specify the boundary between organs by positive and negative feedback regulation of each other’s expression. To further fine tune this regulatory pathway, *STM* mobilization in the meristem regulates spatial expression of the CUC genes to affect organ boundary specification (Balkunde et al., 2017).

Around 25 years ago, Clark et al. (1996) used a genetic approach to suggest that *STM* and *CLV* play opposite and perhaps competitive roles in regulation of meristem cell differentiation. The *stm-1* mutant was previously shown to lack an embryonic shoot meristem (Barton and Poethig, 1993). By comparing the strong *stm-1* allele with the intermediate *stm-2* allele, Clark et al. (1996) observed a variable number of densely stained cells, corresponding to the meristematic cells of the embryonic shoot meristem, only in *stm-2* and not in *stm-1*. These cells were present at relatively similar positions as the wild-type shoot meristem. In *stm-1* seedlings, the region between the cotyledons and the hypocotyl was enlarged relative to wild-type. This enlarged region corresponds to an area of fused cotyledon petioles and not the hypocotyl (Clark et al., 1996). The petiole fusion phenotype is confirmed by internal vascular structures. Early during organogenesis, vascular tissues are differentiated. During leaf development, large primary veins first form, then slightly narrower secondary veins branch from the primary veins, followed by tertiary and quaternary veins. The vein in the hypocotyl branches to create veins into the cotyledons and the true leaves via their respective petioles. This vein branch point is located at the junction of the hypocotyl and the base of the petioles. A certain percentage of *stm-1* plants were able to develop leaves from the axils of the cotyledons, but most of the others died without producing leaves. However, the intermediate *stm-2* plants produced visible leaves and later extensive shoot development. This suggests that other factors, such as other KNOX transcription factors, in the *stm-1* plants are sometimes capable of inducing leaf production. This is supported by the fact that the *stm* mutant sometimes produces abnormal axillary branches (Hake et al. 2004; Scofield and Murray 2006; Hay and Tsiantis 2010; Balkunde et al. 2017).

Clark et al. (1996) also studied how the *clv* mutation can rescue the *stm* phenotype. Even heterozygous *clv1* or *clv3* was able to increase the frequency of leaf development in *stm-1* plants. However, the *clv* mutations do not appear to restore the formation of embryonic meristems in *stm* plants. The *clv stm* mutants have enlarged post-embryonic shoot meristem as compared to wild-type plants. Analysis of floral development of *clv stm-1* plants revealed that *clv* mutations restored floral meristem activity levels of *stm-1* similar level to that of *stm-2* flowers. On the other hand, *stm* dominantly to partially suppressed the *clv3* homozygous phenotype during flower development. In summary, *clv* mutation in the *stm* background caused post-embryonic leaf and floral development. Also, *stm* dominantly suppressed *clv1* semi-dominance based on the carpel development phenotype. Clark et al., 1996 concluded that “a wild-type level of CLV activity is required for the lack of post-embryonic meristem development observed in *stm* mutants and a wild-type level of STM activity is required for the meristem over-proliferation phenotype of *clv* mutants”.

*Arabidopsis* encodes four Class I KNOX transcription factors, namely *STM, KNAT1/BREVIPEDICELLUS* (*BP*), *KNAT6*, and *KNAT2* (Hake et al., 2004). *BP/KNAT1* is involved in internode development (Douglas et al., 2002; Venglat et al., 2002; Smith and Hake, 2003) and contributes to SAM maintenance in conjunction with *STM* (Byrne et al., 2002). *KNAT2* has not yet been shown to have a direct effect of meristem development or activity. *KNAT6* contributes to SAM maintenance and boundary establishment during embryogenesis (Belles-Boix et al., 2006). Using the weak *stm* allele, *knat6-1 stm-2* mutant plants lack a SAM and fail to produce primordia. While both the strong *stm-1* and intermediate *stm-2* show fusion of the petiole of the cotyledons (Clark et al., 1996; Long et al.,1996), the fusion is more pronounced in *knat6-1 stm-2*, extending to the lamina as a result of significantly reduced *CUC3* expression (Belles-Boix et al., 2006). Additionally, *knat6-1 stm-2* plants do not develop further, unlike *stm-2* mutants that continue to make organs post-embryonically (Belles-Boix et al., 2006). In summary, the removal of *KNAT6* gene function eliminates any residual meristematic activity of the *stm* mutants (Belles-Boix et al., 2006). Interestingly, STM regulates KNAT6 activity in the SAM, thus suggesting an additive function of KNOX genes during SAM development.

The central role of STM is integrated into the wider framework of developmental regulation. STM functions via a complex Gene Regulatory Network (GRN) to regulate multiple facets of plant development, such as control of organ formation and differentiation, regulation of pluripotency and phyllotaxis, and the establishment of meristem-organ boundary zones in addition to regulating genes involved in cell wall modification and hormone synthesis and response (Scofield et al., 2018).

Previous studies have explored the interdependent relationship of *STM* and *CLV* during SAM development without taking into consideration that there are other KNOX factors such as KNAT6 in *Arabidopsis*. STM cannot be the only gene responsible for producing the SAM, because loss of the STM protein in the strong *stm-1* mutant does not stop formation of leaves, shoots or flowers in a *clv stm* mutant background. The epistatic behavior between *STM* and *CLV* genes are unclear. In this study, we explored the relationship between *STM* and *CLV3* using strong *stm-1* allele and *KNAT6* to deplete the plants of an additional KNOX factor that has also been shown to have a role, with *STM*, in SAM formation. Our data suggests that *STM* and *KNAT6* are epistatic to *CLV3* in embryonic meristem formation and in organ separation.

## Materials and Methods

### Plant Growth Conditions

All plants used in this study were grown at 22 °C under long-day (16 hours) illumination. Seeds were germinated on solid sterile medium containing 4.3 g MS Medium and 0.5 g MES buffer per liter at pH 6.5, adjusted with 1 M KOH, and 0.8 g phytagar per liter. Soil-grown plants were seeded in ProMix PGX soil mix, and Osmocote slow-release fertilizer was added to the flats as directed by manufacturer.

### Plant Genetics

Both *stm-1* and *clv3-1* are in the *Landsberg erecta* (Ler) background, whereas *knat6-1* is in the Col-0 background. *stm-1* and *clv3-1* mutations were phenotypically genotyped using Mendelian inheritance ratios of the meristemless phenotype and enlarged inflorescence meristem phenotypes, respectively. *knat6-1* is a T-DNA insertion line and was genotyped by PCR using Lba1 T-DNA primer and *KNAT6*-specific primer (primer sequences are listed in Supplementary Table 1).

To generate *stm-1 clv3-1 knat6-1* triple mutants, plants homozygous for *clv3-1* were crossed to plants heterozygous for *stm-1*. The shoot meristemless phenotype segregated in the F2 progeny in the expected 1:16 ratio. F2 plants homozygous for *clv3-1* and heterozygous for *stm-1* were selected based on segregating meristemless and enlarged meristem phenotypes. This line was then crossed to plants homozygous for *knat6-1*. A novel triple mutant phenotype segregated in the F2 progeny that resembled the published *knat6-1 stm-2* phenotype (Belles-Boix et al., 2006) in addition to the *stm-1* meristemless phenotype.

### Microscopy

Eight days after germination (DAG), seedlings were imaged using a Leica compound microscope, brightfield filter and a 20X objective. For Nomarski interference contrast (NIC) microscopy, 8-DAG seedlings were cleared in 7:1 ethanol:acetic acid, treated for 30 min with 1 N potassium hydroxide, rinsed in water, and mounted in Hoyer’s medium. NIC images were obtained using a Leica compound microscope equipped with Nomarski optics and a 20X objective. For Scanning Electron Microscopy (SEM) imaging, 21-DAG fresh tissue was directly imaged using FEI Quanta 200 Environmental SEM.

### Quantitative analysis of seedling phenotype

We used the formula for calculating the area of an ellipse (π × semi-major axis × semi-minor axis) to quantify the area of the inflorescence (Heath 1897) The radius across the inflorescence was measured twice, at a 90° angle, with the measurements labeled as r1 and r2. The formula Inflorescence area = π × r1 × r2 was used to calculate the size of the inflorescence. Student’s *t*-test was used to confirm statistically significant differences.

### Real-time PCR

RNA was extracted using RNeasy kits from Qiagen. First-strand cDNA was made from 2 μg of RNA using the Tetro cDNA synthesis kit with DNase treatment according to manufacturer’s instructions (Bioline). cDNA was diluted 1 to 5 into RNAse free water to a total of 100 μl. The diluted cDNA (2 μl) was used for qRT-PCR. PCR was done using gene-specific primers in technical triplicates on a LightCycler 480 system using the Sensifast SYBR Master mix (Bioline). The ratio of experimental target mRNA to Actin control for each sample was calculated by Applied Biosystems software. An average and standard error of the mean for the three biological replicates and standard deviation were calculated in Microsoft Excel. Primers are listed in Supplementary Table 1.

## Results

### Construction of *stm clv3 knat6* triple mutant

To understand the epistatic relationship between KNOX and CLV genes, we generated double and triple mutant plants using *stm-1/+, clv3-1*, and *knat6-1* mutant alleles. These alleles have already been described. We crossed *stm-1/+* and *clv3-1/-* mutant plants to generate *stm-1/+ clv3-1* double mutants. The segregating F1 population of *stm-1/+ clv3-1/+* were self-pollinated. In the F2 generation, we confirmed the *stm-1* meristemless phenotype at the seedling stage among the independent lines. To confirm the presence of *clv3-1*, the plants were grown to flowering stage and checked for the enlarged inflorescence meristem phenotype. Lines that showed the *clv3-1* mutant phenotype in at least 40 plants and also a segregating 1:4 *stm-1* phenotype were selected. In our assumption, these plants were *stm1-/+ clv3-1/-*. The *stm-1/+ clv3/-* F3 plants were grown and crossed to *knat6-1/-* homozygous mutant plants to generate higher order mutants. We followed the same process to find *stm-1* and *clv3-1* genotypes as described above. Since *knat6-1* is a T-DNA insertion allele, we used PCR based genotyping to identify *knat6-1/-* alleles in the *stm-1/+ clv3/-* plants. We repeated this process for an additional two generations and identified *stm-1/+ knat6-1/-* double, *clv3-1/- knat6-1/-* double, and the *stm-1/+ clv3/- knat6/-* triple mutant plants from the population. For simplicity, the triple mutant line is labeled as *stm-1 clv3-1 knat6-1* triple mutant for the rest of the paper.

### Characterization of *stm-1 clv3-1 knat6-1* triple mutant plants

STM is necessary for both development and maintenance of the shoot apical meristem (SAM). Wild-type Col-0 and Ler have normal SAM, and thus, produce true leaves (Figs 1A and 1B) and floral buds (Fig 2A). On the other hand, *stm-1* mutants lack a SAM and fail to produce true leaves (Fig 1C and 2B). Also, in the *stm-1* seedlings, the region between the cotyledons and the hypocotyl is enlarged relative to wild-type. This enlarged region corresponds to an area of fused cotyledon petioles and not the hypocotyl. (Fig 1C and 2B). Nomarksi Interference Microscopy (NIC) was performed to further observe the organ fusion by visualizing the internal structures as well as cell types within the fused region (Fig 1 bottom panel sets). The NIC images of the vasculature pattern of the wild type and *stm-1* mutant provide additional information that explains the organ-fusion phenotype. In our observations, wild-type Col and Ler plants showed normal branching of the hypocotyl vein into the cotyledon and true leaf petioles below the SAM, where new leaves are formed (Figs 1A and B bottom panels, arrowhead). In contrast, the branch point of the hypocotyl vein in the *stm-1* mutant was lower than the place of petiole emergence, suggesting the petioles are fused in that region (Fig 1C bottom panel).

**Figure 1:**
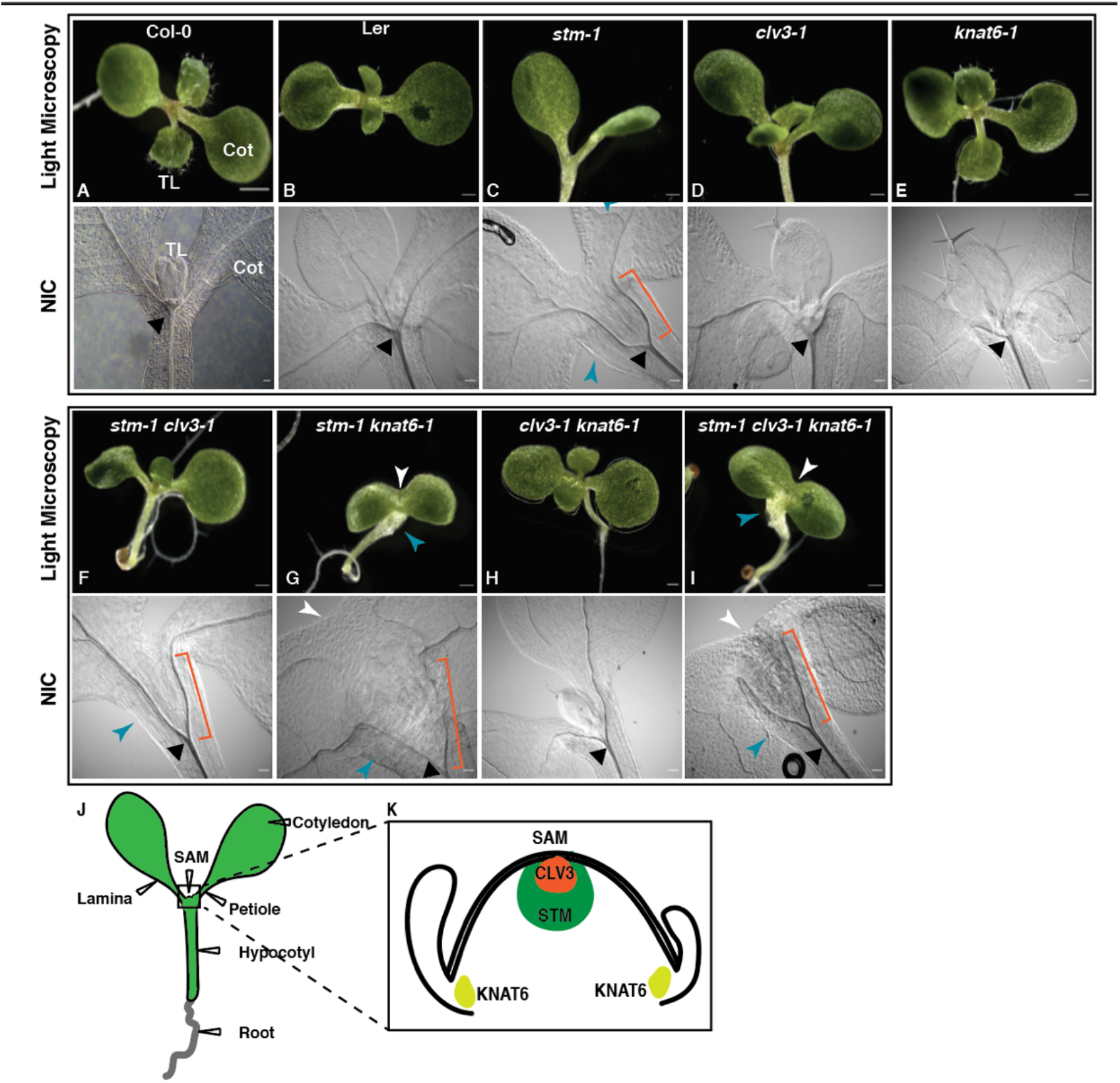
Seedling phenotypes (8 DAG) under light and Nomarski Interference Contrast (NIC) microscopy. **A** and **B**: Col-0 (**A**) and Ler **(B)** plant with true leaves due to a functional shoot apical meristem (SAM). Cot, cotyledon; TL, True Leaf. **C**: *stm-1* mutant plants lacked SAM and true leaves, and had fused petioles (blue arrowhead). **D** and **E**: *clv3-1* (**D**) and *knat6-1* (**E**) single mutants with functional meristem and true leaves. **F**: *stm-1 clv3-1* double mutants produced true leaves at a lower frequency as compared to *clv3-1*.**G**: *stm-1 knat6-1* double mutants lacked SAM and showed fusion of cotyledons (white arrowhead) and petioles (blue arrowhead). **H**: *clv3-1 knat6-1* plants have functional meristem and produce true leaves. **I**: *stm-1 clv3-1 knat6-1* triple mutants lacked SAM and showed fusion of cotyledons (white arrowhead) and petioles (blue arrowhead Bottom Panels: NIC microscopy images of wild-type Col-0 (**A**) and Ler (**B**), *clv3-1* (**D**), *knat6-1* (**E**), and *clv3-1 knat6-1* (**H**) with a hypocotyl vein branch point (black triangle) below the SAM region. NIC microscopy images of *stm-1* (**C**), *stm-1 clv3-1* (**F**), *stm-1 knat6-1* (**G**), and *stm-1 clv3-1 knat6-1* (**I**) mutants with a lower hypocotyl vein branch point (black triangle) more distant from the SAM region. The orange bracket shows the extent of petiole fusion using NIC microscopy. **J.** A schematic diagram showing various parts of a seedling. **K.** An inset of the SAM region showing published expression domains of *STM, KNAT6*, and *CLV3*. Scale bars are 100um each.

**Figure 2:**
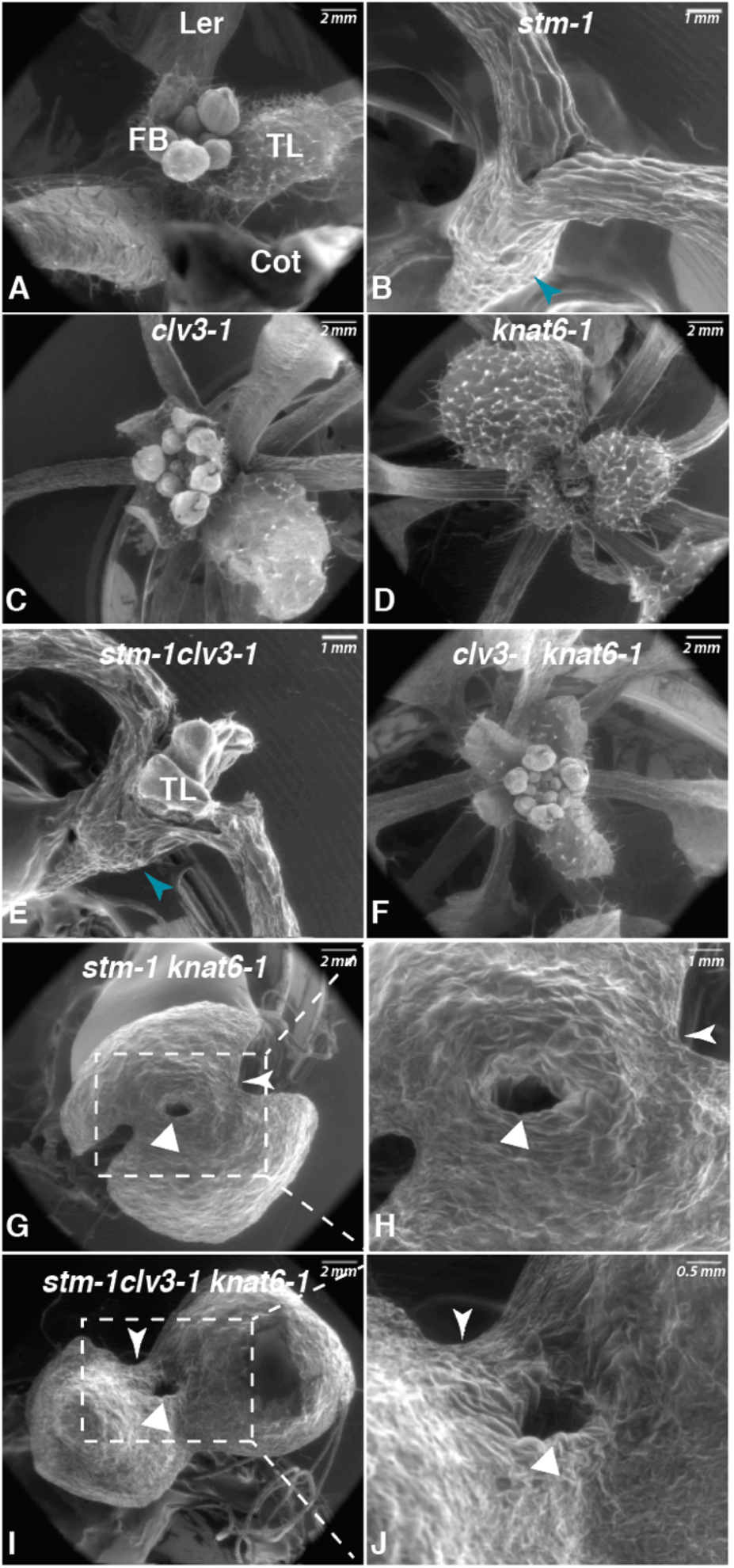
Seedling phenotype of 15 dag plants under Scanning Electron Microscope (SEM) **A**: Ler plants with normal SAM and true leaves (TL) and floral buds (FB). Cot, cotyledon. **B**: *stm-1* single mutant plants without SAM or TL but with fused petiole (blue arrowhead). **C**: *clv3-1* mutants with functional SAM and true leaves and floral buds. **D**: *knat6-1* plants with functional SAM and TL. **E**: *stm-1 clv3-1* double mutants with fused petioles (blue arrowhead) and occasionally produce true leaves from ectopic meristems. **F**: *clv3-1 knat6-1* double mutant with normal SAM, TL, and FB. **G** and **H**: *stm-1 knat6* double mutants without SAM and with fused cotyledon boundaries (white arrowhead). The lack of tissue at the presumed site of SAM is evident (white triangle). **H** is a zoomed-in image of **G**. **I** and **J**: *stm-1 clv3-1 knat6-1* triple mutants without SAM and with fused cotyledon boundaries. The presumed site of SAM lacked tissue (white triangle). **J** is a zoomed-in image of **I**. Scale bars are individually labeled.

C*LV3* is known to dampen stem cell proliferation by negatively regulating WUS expression (Perales and Reddy 2012; Nimchuk, et al 2015). We observed that *clv3-1* mutant seedlings have functional meristem and, thus, give rise to true leaves and floral buds similar to wildtype (Figs 1D and 2C). The hypocotyl vein node in *clv3-1* is placed below the SAM region the same as wildtype (Fig 1D bottom panel). The majority of the *stm-1 clv3-1* double mutants lacked an embryonic SAM; however, some of the plants ultimately produced true leaves (Figs 1F and 2E). This phenotype has been reported earlier by Clark et al, 1996. We further observed that *stm-1 clv3-1* plants also had fused cotyledon petioles and a lower hypocotyl vein branch point, similar to the *stm-1* mutant (Fig 1F bottom panel).

STM belongs to the KNOX family of transcription factors, which includes the closely related homologue KNAT6. To explore the genetic interaction among those genes and CLV3, we generated triple mutants of *stm-1 knat6-1 clv3* and compared them to the corresponding single and double mutants. Unlike *stm-1*, the *knat6-1* single mutant has a functional SAM and produces true leaves similar to wild-type plants (Figs 1E and 2D). The hypocotyl vein branch point in *knat6-1* is placed below the SAM region, the same as wild-type plants (Fig 1E bottom panel). Interestingly, *stm-1 knat6-1* also lacked a SAM, and the fusion of the cotyledon petiole is extended to include fusion of the cotyledons and lamina as well (Figs 1G and 2G). The hypocotyl vein branch point in the *stm-1 knat6-1* mutant is in a lower position, the same as *stm-1* (Fig 1G bottom panel). The fusion of cotyledons and lamina in *stm-1 knat6-1* (Fig 1G) is similar to the *stm-2 knat6-1* mutant (Belles-Boix et al., 2006). Notably, Scanning Electron Microscopy (SEM) revealed an exceptional SAM phenotype in these mutants. We observed an absence of tissue in place of a SAM in the *stm-1 knat6-1* mutants (Figs 2G-H). The presumed SAM region is hollow in *stm-1 knat6-1* mutants (Figs 2G-H). This is a first report of such phenotype. *clv3-1 knat6-1* plants have functional SAM and produce true leaves, similar to the *clv3-1* and *knat6-1* single mutants (Figs 1H and 2F). Additionally, *clv3-1 knat6-1* mutant plants show normal hypocotyl vein branch point placement, below the SAM, the same as wildtype (Fig 1H bottom panel). This single and double mutant analysis suggests that *clv3-1* is epistatic to *stm-1*, while *stm-1* and *knat6-1* have additive roles in the formation and maintenance of SAM. However, in the *stm-1 clv3-1* background, the *KNAT6* gene is functional and could be leading the postembryonic SAM function. In order to study the effects of absence of both *STM* and *KNAT6* genes in the *clv3-1* background, we compared the SAM in the *stm-1 clv3-1 knat6-1* triple mutants with the respective single and double mutants. The *stm-1 clv3-1 knat6-1* triple mutants were similar to the *stm-1 knat6-1* double mutants, as *stm-1 clv3-1 knat6-1* also lacked SAM and showed severe fusion of cotyledon, lamina, and petiole phenotype (Figs 1I, 2I, and 2J). The *stm-1 clv3-1 knat6-1* mutants had a lower hypocotyl vein branch point, the same as *stm-1* (Fig 1I bottom panel). SEM revealed the triple mutant also lacked tissue in place of SAM (Figs 2 I-J), much like *stm-1 knat6-1* (Figs 2G-H). The triple mutant phenotype analysis suggests that *stm* is epistatic to *clv3* in the formation of embryonic SAM contrary to previously published data by Clark et al (1996).

### Phenotypic variability within the segregating population of mutant alleles

Unlike wild-type and the intermediate *stm-2* allele (Clark et al., 1996), the strong *stm-1* allele gives rise to an enlarged region between the cotyledons and the hypocotyl. This enlarged region corresponds to an area of fused cotyledon petioles and not the hypocotyl (as described above and in Fig 1). This fused cotyledon phenotype is enhanced in the *knat6-1 stm-2* seedlings, where the fusion of cotyledons is extended to the lamina (Belles-Boix et al., 2006). We also observed the enhanced fusion of the cotyledon, lamina, and petioles in the *stm-1 clv3-1 knat6-1* triple mutant seedlings (Fig 2).

In the mutant lines we generated, the *stm-1* allele is heterozygous to allow formation of leaves, flowers, and seeds in the following generations. Thus, we observed variability of the meristemless phenotype at the seedling stage in the segregating population of around 200-400 plants (See supplementary Table 2 for N-values). We categorized the phenotypes into 3 groups: Normal SAM (M); No SAM (m); and No SAM + Fused cotyledon/lamina/petiole (m+) (Fig 3). Plants with “Normal SAM (M)” produce leaves normally (blue in Fig 3), plants with “No SAM (m)” lacked a meristem and did not produce leaves, much like *stm-1* (purple); and plants with “No SAM + Fused cotyledon/lamina/petiole (m+)” lacked a meristem and showed enhanced fusion of cotyledon, lamina, and petiole (yellow). The entire populations of wildtype Col-0 and Ler were phenotype M. In the segregating *stm-1/+* population, 28% plants had the m phenotype. The *clv3-1, knat6-1*, and *clv3-1 knat6-1* mutants were 100% phenotype M. In the *stm-1/+ knat6/-* double mutant population, which was segregating for *stm1* but not *knat6-1*, 74% plants had the M phenotype, about 1% had the m phenotype, and 25% were m+ phenotype. The *stm-1/+ clv3-1* double mutant population was 78.5% M, 21% m, and 0.5% m+ phenotype. The *stm-1/+ clv3-1 knat6-1* triple mutant population was 73% M, about 0.2% m, and 26% m+. This data suggested that the extent of organ fusion was not affected by the *clv3* mutation in the *stm-1 knat6-1* background.

**Figure 3:**
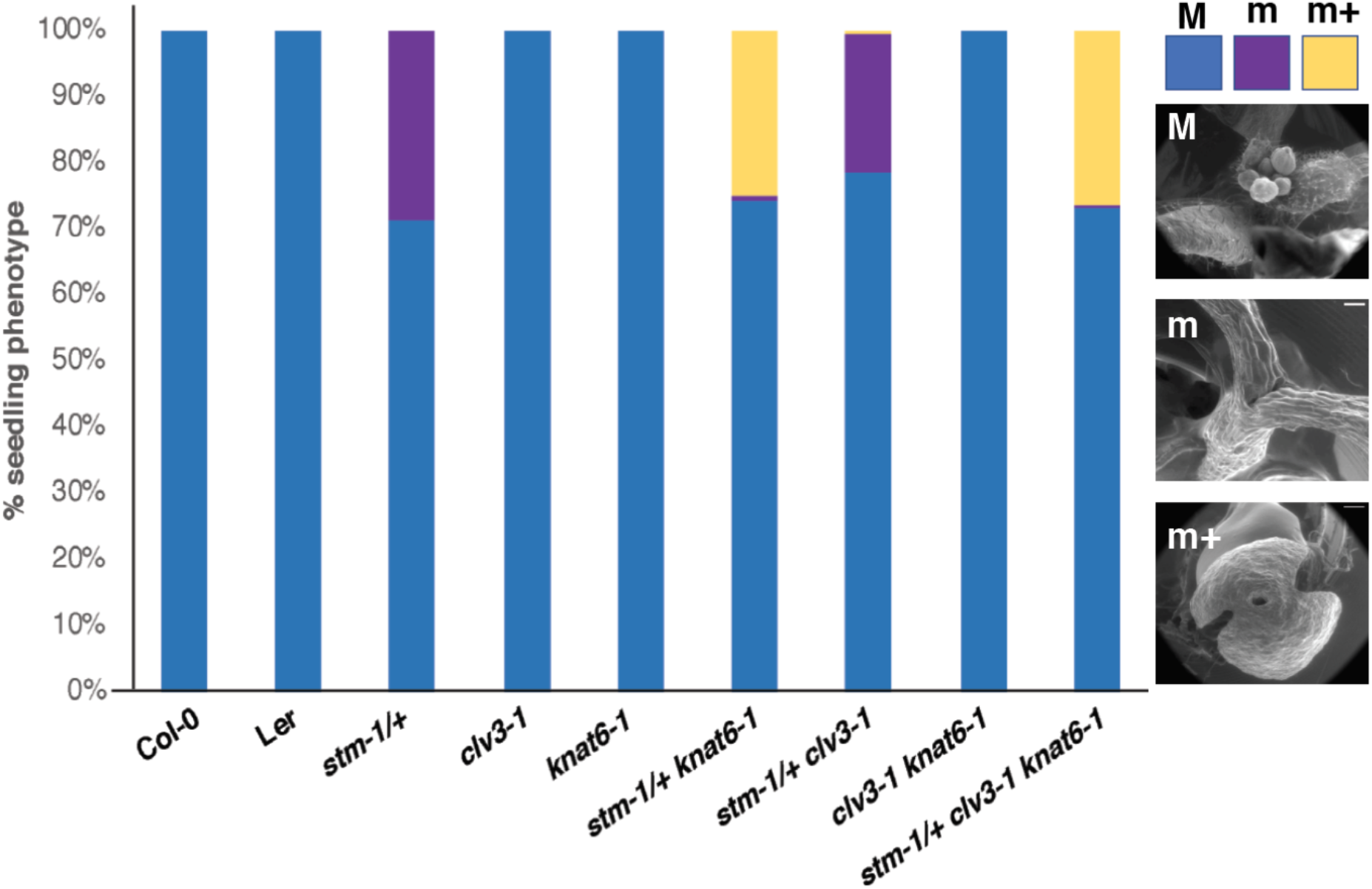
Quantification of SAM phenotype in seedlings of nine genotypes. Phenotypes were assessed 10 Dag. M represents a normal SAM, m represents lack of SAM, and m+ represents lack of SAM in addition to fusion of cotyledons and petioles. Inset images show the respective phenotypes. The *stm-1* mutation was the only genotype segregating. N-values are listed in Supplementary Table 2.

### *CLV3* is epistatic to both *stm* and *knat6* during inflorescence meristem development

CLV3 negatively regulates inflorescence meristem size, thus, *clv3* mutants have an enlarged inflorescence meristem resulting in a bigger inflorescence diameter. The *stm* mutation partially rescues the enlarged floral meristem in *clv3* mutants (Clark et al., 1996). To investigate the effects of the KNOX genes on inflorescence size in *clv3-1* mutants, we quantified the inflorescence area by using the formula for calculating the area of an ellipse (π × semi-major axis × semi-minor axis). We measured the radius across the inflorescence at two points at a 90° angle to each other (r1 and r2) and used the formula π × r1 × r2 to calculate the area of the inflorescence. (Fig 4I). We compared the inflorescence area of the double and triple mutants to *clv3-1* single mutants, which possess the largest inflorescence area among the tested genotypes (Fig 4D, J and K). Wild-types Col-0 (Fig 4A) and Ler (Fig 4B) showed normal inflorescence areas, ranging from 10 to 70 mm^2^/cm plant height. The inflorescence areas of *knat6-1* (Fig 4C) and *stm-1* (Fig 4E) were similar to Col-0 and Ler, respectively. As expected, the *stm-1/+ clv3* plants have a smaller inflorescence area as compared to *clv3-1* plants (Fig 4F, J, and K). Interestingly, *knat6-1* reduced the inflorescence size in the *clv3-1* background by 65%, as seen in the *clv3-1 knat6-1* double mutant (Fig 4G, J, and K). The *knat6-1* was statistically similar to *stm-1 knat6-1* inflorescence area (Fig 4K). However, absence of both *STM* and *KNAT-6* in the *stm-1 clv3-1 knat6-1* triple mutant resulted in a 52% reduction in the inflorescence area as compared to *clv3-1* plants (Fig 4H, J, and K). Thus, *stm* and *knat6* mutations have an additive effect in regulating *clv3* inflorescence size. This suggests that *CLV3* is epistatic to both *STM* and *KNAT6* during inflorescence meristem development.

**Figure 4:**
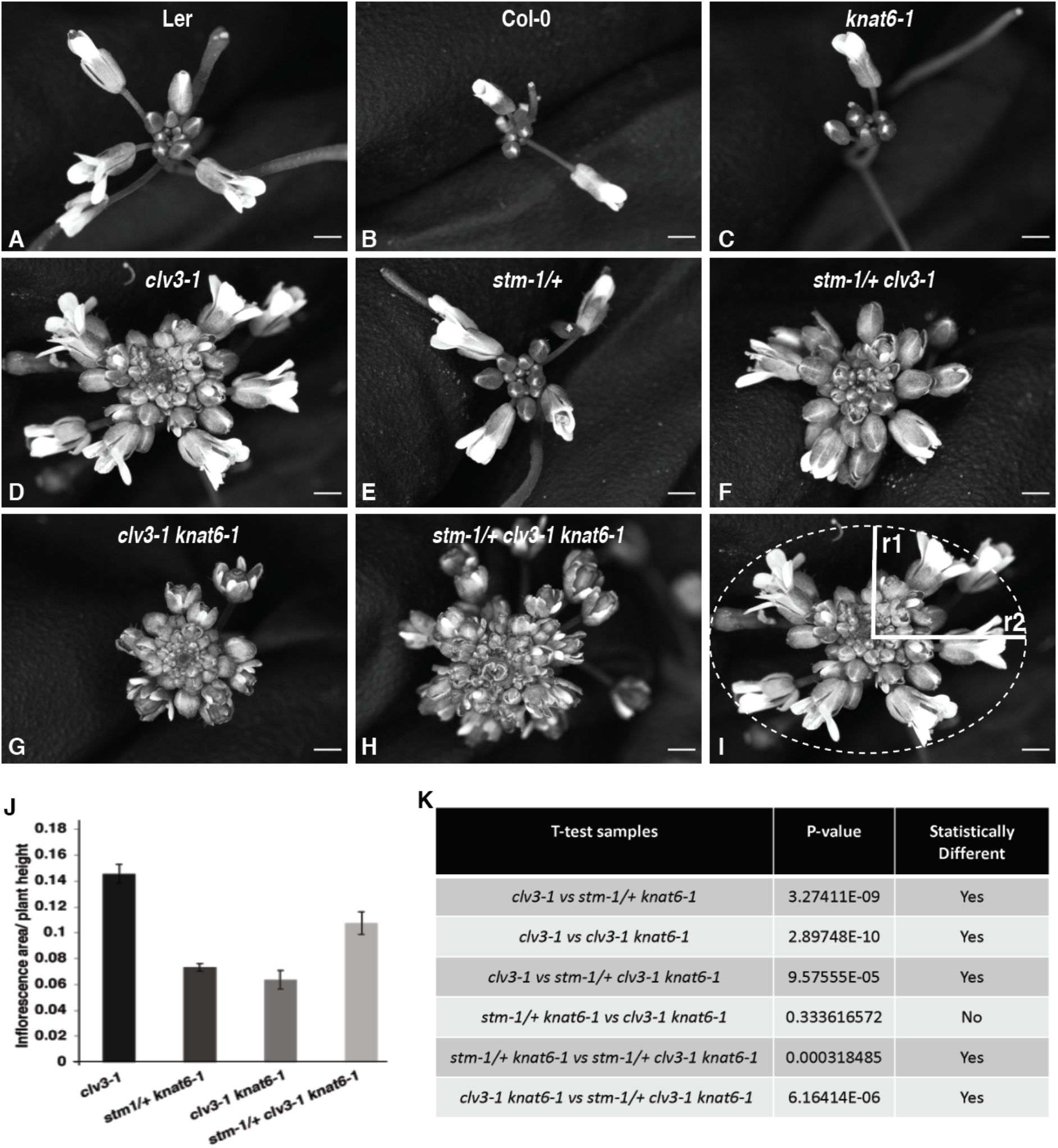
Quantification of inflorescence size in adult plants. **A-H:**Brightfield images of inflorescences from the top of the adult plants: Ler (**A**), Col-0 (**B**), *knat6-1* (**C**), *clv3-1* (**D**), *stm-1/+* (**E**), *stm-1/+ clv3-1* (**F**), *clv3-1 knat6-1* (**G**), and *stm-1/+ clv3-1 knat6-1* (**H**). **I, J**: Quantification of inflorescence sizes. A representative image showing the radii measured at perpendicular angles (I). Inflorescence area was measured using the formula Inflorescence size = π × r1 × r2, where r1 and r2 are radii measured at at 90°to each other. The inflorescence area was normalized with the height of the plant. The genotypes were compared against *clv3-1* (**J**). **K**: Pairwise Student’s *t*-test results (*p*-value) to confirm statistical significance in difference. N-values are as follows: *clv3-1* = 24; *stm1/+ knat6-1* = 46; *clv3-1 knat6-1* = 53; *stm1/+ clv3-1 knat6-1* = 49.

### Expression of *CLAVATA3* gene in response to *STM* activation

STM and CLV3 have been shown to have biologically antagonistic functions. *STM* promotes meristem formation, while *CLV3* represses meristem size. We employed glucocorticoid receptor-induced gene expression system to test if *STM* activation has an effect on *CLV3* gene expression. We generated 35Sp::GR-STM transgenic plants in Col-0 background, and activated the transgene with Dexamethasone (DEX) treatment. The 35Sp::GR-STM transgenic plants grown on 50 μM Dexamethasone were smaller than Col-0 and produce ectopic meristematic cells at the shoot apex (Fig 5A). We tested STM induction by growing 35Sp::GR-STM transgenic plants in liquid MS media and treating them with 50 μM Dexamethasone for 30 min. With DEX treatment, STM gene expression was 1.6-fold higher than the untreated samples (Fig 5B). Interestingly, *CLV3* gene expression was repressed with DEX treatment (Fig 5B). This result suggested that STM gene activation and of *CL3V* gene repression are correlated.

**Figure 5:**
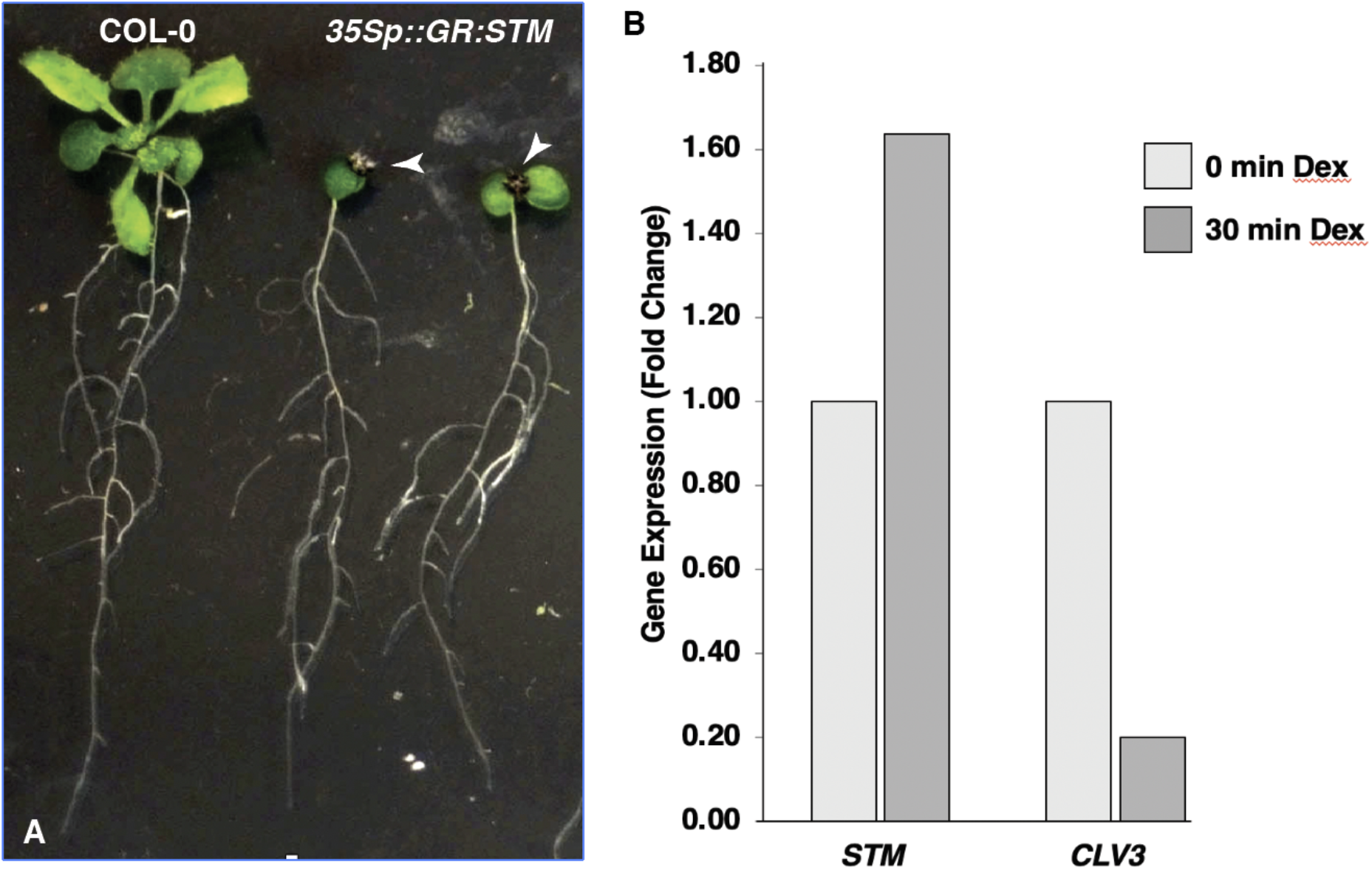
Relative gene expression levels of STM and CLV3. **(A)** Seedling phenotypes of Col-0 and 35Sp::GR-STM lines grown on MS media + 50 μM Dexamethasone (DEX). The meristem region in the 35Sp::GR-STM plants had small purple leaves and ectopic meristematic cells (white arrowhead). **(B)** Relative gene expression levels of STM and CLV3 in 35Sp::GR-STM plants untreated or treated with 50 μM Dex for 30 min. *STM* was induced upon Dex treatment, and *CLV3* expression was reduced upon Dex treatment.

## Discussion

Both animals and plants rely on the generation of new cells for growth and development. In plants, the shoot apical meristem (SAM) consists of a population of self-renewing, or stem, cells, and provides daughter cells for initiation and development of aboveground organs. However, the underlying mechanisms regulating the size of the SAM remain largely unclear.

It is essential to acknowledge the range of SAM functions that are coordinated through the *SHOOTMERITEMLESS* (*STM*) Gene Regulatory Network. This places STM in a central role during development. STM, a member of the KNOX family of transcription factors, coordinates organ formation and differentiation, regulation of pluripotency and phyllotaxis, the establishment of meristem-organ boundary zones, in addition to regulating genes involved in cell wall modification and hormone synthesis and response (Scofield et al., 2018).

*stm-1* mutants fail to form an embryonic SAM. However, if grown for an extended period of time, the *stm-1* mutant produce ectopic leaves at a low frequency, suggesting post-embryonic meristem development and activity. There are three other members in the KNOX transcription factor family, namely *KNAT1* or *BREVIPEDICELLUS* (*BP*), *KNAT6*,and *KNAT2*. Presumably, these KNOX proteins are present in the *stm-1* background and could hypothetically be leading the post-embryonic meristem development. Here we show that STM and KNAT6 are necessary to the formation of embryonic and post-embryonic meristem cells. In the absence of *stm* and *knat6*, plants lack SAM and form fused cotyledons, at both the lamina and petiole. The *stm-1 knat6-1* double mutants did not induce ectopic leaves when allowed to grow for an extended time (data not shown). Earlier, Belles-Boix et al. (2006) showed a similar phenotype in the double mutant of *knat6* and the weaker *stm-2* allele. We studied the stronger *stm-1* allele in this study to ensure complete reduction of STM protein.

CLAVATA (CLV) and STM have been known to have opposite functions, because *stm* mutants fail to form the undifferentiated cells of the shoot meristem during embryonic development, while *clv1* and *clv3* mutants accumulate excess undifferentiated cells in the shoot and floral meristem. Previously, it was reported that *clv1* and *clv3* mutations partially suppressed the *stm-1* and *stm-2* phenotypes in a dominant manner (Clark et al. 1996). It is important to note that *stm-1* is a strong allele, whereas *stm-2* is a weak allele. While the *clv3 stm-2* double mutant lacks the embryonic SAM, it still exhibited greatly enhanced postembryonic shoot and floral meristem development. This suggested that CLV3 is epistatic to STM in post embryonic meristem development. Interestingly, it was shown that *stm* mutation dominantly suppressed the *clv* phenotype at the inflorescence meristem. Clark et al (1996) concluded that the *stm* phenotype is sensitive to the levels of CLV activity, while the *clv* phenotype is sensitive to the level of STM activity.

Contrary to previously published conclusions, our results showed that STM along with KNAT6 are epistatic to CLAVATA3. Using a *stm-1 clv3-1 knat6-1* triple mutant, we showed that *STM* and *KNAT6* act upstream of *CLV3* during embryonic and post-embryonic meristem development. These results indicate that the previous studies did not consider other (unknown) members of the KNOX family. In our experiments, the *stm-1 clv3-1 knat6-1* triple mutant plants showed similar phenotypes (lack of embryonic meristem and extended organ fusion) as the *stm-1 knat6-1* double mutant.

The fusion of the cotyledons observed in the *stm-1 knat6-1* double and *stm-1 clv3-1 knat6-1* triple mutants suggests that the *CUP-SHAPED COTYLEDON* (*CUC*) genes are involved with *STM* and *KNAT6*. The *CUC* gene family plays a crucial role in the establishment of the SAM and in organ separation during embryogenesis (Aida et al., 1999). The majority of the *cuc1-1, cuc2*, and *cuc3-2* single mutants have separate cotyledons, however, double mutant combinations of *CUC1, CUC2*, or *CUC3* lead to cup-shaped seedlings (Aida et al., 1999). This suggests redundancy among the *CUC* genes. Belles-Boix et al. (2006) studied interaction of *STM, KNAT6*, and *CUC* genes in detail. In their study, they found that *CUC3* expression is downregulated in *knat6-1 stm-2* embryos and absent in the double mutant seedlings as compared to *knat6* single mutant. This was supported by the phenotype of the *knat6-1 stm-2 cuc3-2* triple mutant, as the *cuc3* mutation enhanced the fusion present in the *knat6-1 stm-2* phenotype. The fusions were less severe in the absence of *CUC1* or *CUC2* (as compared to *CUC3*) in the *stm-2 knat6-1* background, suggesting residual *CUC1* or *CUC2* activity in the respective triple mutants (Belles-Boix et al., 2006). For future studies, it will be interesting to see effects of the *cuc* gene mutations in the *stm-1 knat6-1* and *stm-1 clv3-1 knat6-1* backgrounds, since *stm-1* is a strong allele as compared to *stm-2*. In the strong *stm-1* background, we hypothesize that the CUC proteins will be depleted and that severe fusion of cotyledons in the *stm-1 knat6-1 cuc1/2/3* mutants will be seen.

We also investigated the interaction of *STM, KNAT6*, and *CLV3* at the inflorescence meristem stage. The *clv3* mutants have enlarged floral meristem, as observed by the dense floral buds at the shoot apex of flowering Arabidopsis. As stated above, Clark et al. (1996) showed that the *stm* mutation dominantly suppressed the *clv* inflorescence meristem phenotype. Similarly, we found that *stm-1* and *knat6-1* mutations reduced the *clv3* inflorescence meristem size in the triple mutant, but not to the wild-type level, suggesting that *CLV3* is epistatic to *STM* and *KNAT6* during inflorescence meristem development.

Our gene expression data suggests a correlation between STM activation and depletion of *CLV3* expression, suggesting that STM activation leads to downregulation of CLV3. A recent report by Su et al. (2020) shows that STM protein directly binds to the *CLV3* promoter. The binding of STM to the *CLV3* promoter is required for *CLV3* promoter activity and essential for *CLV3*-mediated stem cell control. This further strengthens our conclusion that STM activation leads to *CLV3* downregulation.

In this study we explored the interaction between the KNOX gene family and *CLAVATA3* during embryonic and post-embryonic meristem development. We found that *STM* together with *KNAT6* are necessary for the formation of meristem cells in *Arabidopsis*. We also found that *CLV3* acts downstream of *STM* and *KNAT6* during embryonic meristem development, but upstream during inflorescence meristem development. We predict that a fine balance of *KNOX* and *CLAVATA* levels leads to optimum meristem formation and activity for growth and development of plants.

## Acknowledgments

We thank Dr. Kathryn M. Barton and the Carnegie Institute for Science for supporting the work at Carnegie Institution of Science, Stanford, CA. We also thank Dr. Véronique Pautot for providing *knat6-1* seeds. This work is partially supported by University of Florida Seed Grant and startup fund.

## Abbreviations

SAM: Shoot Apical meristem
NIC: Nomarksi Interference Microscopy
GR: Glucocorticoid Receptor
Dex: Dexamethasone

## Author contribution

**Nidhi Sharma**, Conceptualization, Investigation, Data curation, Formal Analysis, Methodology, Validation, Visualization, Writing – original draft, Writing – review & editing; **Tie Liu**, Writing – review & editing; **Kathryn M. Barton**, Conceptualization, Funding acquisition, Supervision.

## Data Availability Statement

The data supporting the findings of this study are available from the corresponding author, Tie Liu, upon request.

## Supplementary Data

**Table S1:**
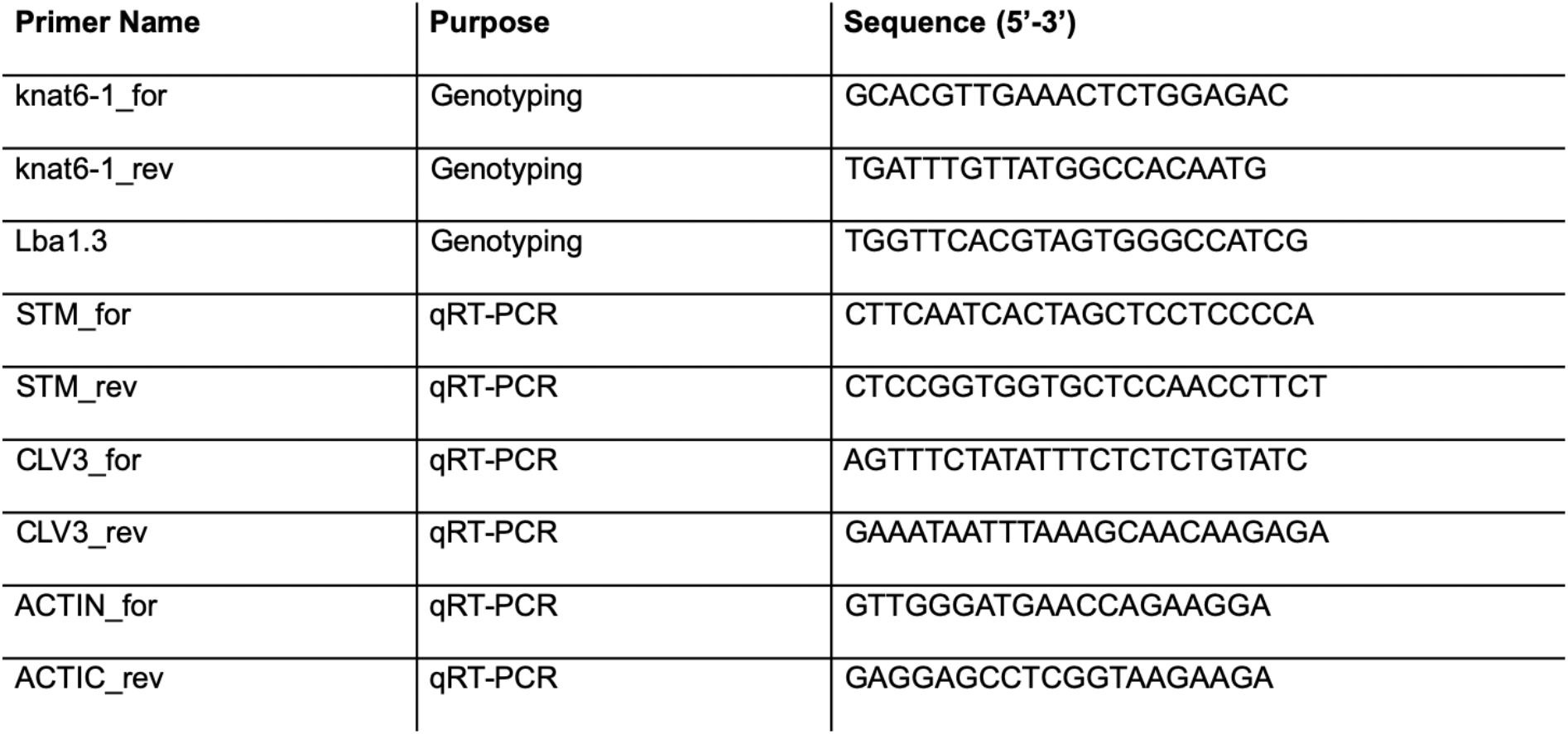
Primer sequences for genotyping and qRT-PCR

**Table S2:**
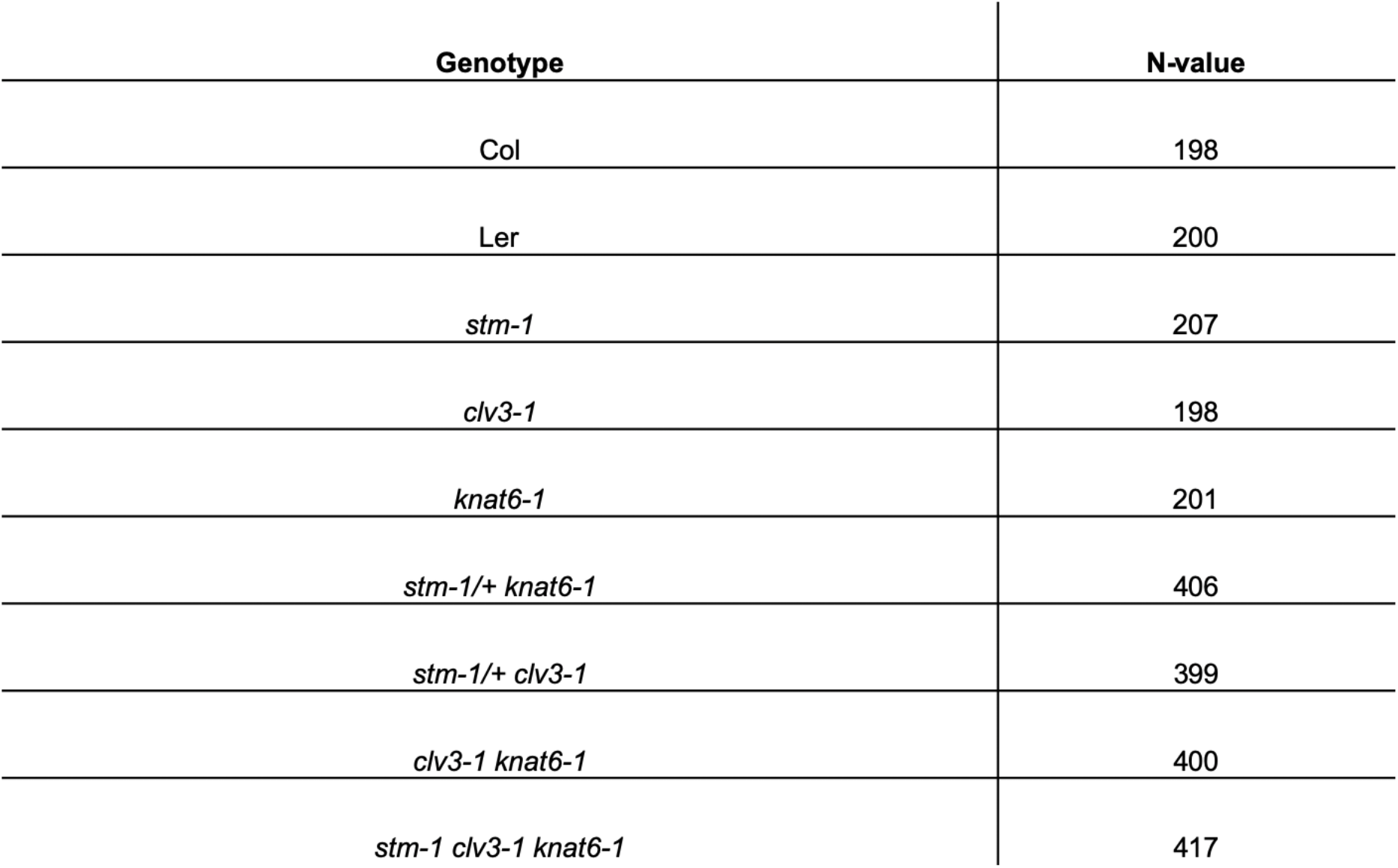
Sample size for meristem phenotype data (Fig 3)

